# Abrupt versus gradual application of pesticides: effects on soil bacterial and fungal communities

**DOI:** 10.1101/2025.03.24.644867

**Authors:** Judith Riedo, Juan F. Dueñas, Susan Mbedi, Sarah Sparmann, Matthias C. Rillig

## Abstract

Pesticides are a major anthropogenic input to the environment and a factor in global change that puts pressure on soil microbial communities. However, the effects of different rates of pesticide application on soils remain poorly understood. This study investigates how abrupt versus gradual pesticide applications influence soil bacterial and fungal communities. Employing high-throughput sequencing, we examined the microbial diversity and community composition in response to ten commonly used pesticides. Bacterial communities exhibited minimal changes across treatments, whereas fungal communities responded strongly to pesticide exposure. Gradual applications reduced the relative abundance of dominant fungal taxa, resulting in an overall increase in community evenness. This effect was particularly pronounced for two herbicides and a triazole fungicide, which induced substantial shifts in fungal composition. Conversely, abrupt pesticide applications resulted in transient disruptions but did not promote the long-term proliferation of rare fungal variants. These findings suggest that prolonged exposure to pesticides exerts strong selective pressures on fungi, potentially altering fundamental soil functions such as nutrient cycling and decomposition. Future research should focus on the long-term responses of soil microbial communities to pesticide application and the cumulative effects of chronic low-dose exposure to provide a more comprehensive understanding of how they shape microbial communities.

**Graphical Abstract:** 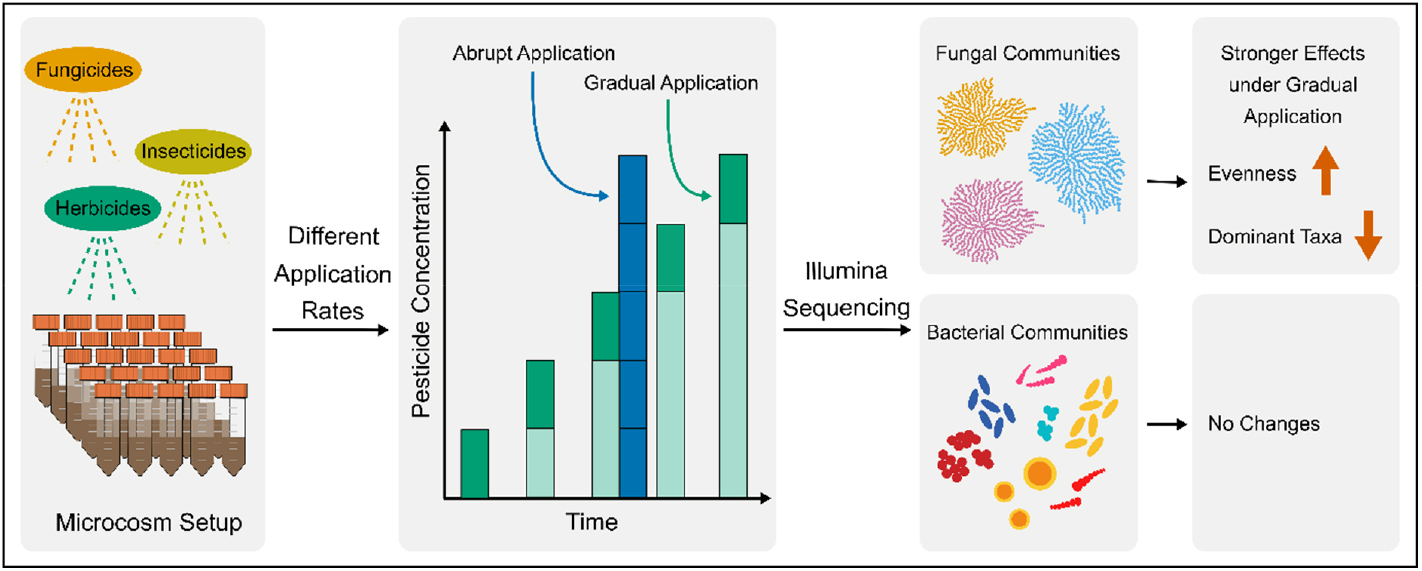

**Highlights:**

- Fungal communities were more responsive to pesticides than bacterial communities
- Herbicides caused strong non-target effects, similar to those of fungicides
- Gradual pesticide applications altered fungal diversity and community composition
- Rare fungal taxa proliferated under gradual exposure, increasing evenness
- Prolonged exposure exerted greater selective pressure than abrupt application

## Introduction

The importance of soil microbial communities to ecosystem functionality is well established. These communities mediate fundamental processes, including nutrient cycling, organic matter decomposition, and soil aggregate formation (Bardgett and van der Putten 2014; Wagg et al. 2014; Schimel 2016). They include diverse bacteria and fungi, which often perform complementary roles. Bacteria are dominant in the early stages of decomposition, while fungi contribute to aggregate stability and nutrient mobilization (Lehmann, Zheng, and Rillig 2017; Fierer 2017). A highly diverse microbial community is fundamental for the resilience of soil systems and can withstand environmental disturbances. Nevertheless, the integrity of these communities is increasingly threatened by various factors of global environmental change, including habitat loss, climate extremes, and chemical pollution, such as pesticides.

Pesticides are a ubiquitous global change factor and constitute one of the largest intentional inputs of hazardous compounds into soil systems, with global usage exceeding 2.5 million tons annually (FAO 2023). Within the European Union, there are over 400 active ingredients that have been registered for use (European Commission 2020). The off-target movement of these substances through drift, runoff, and volatilization following extensive use leads to the accumulation of residues in soils. Although pesticides are considered degradable, these residues can persist in the soil matrix and exert an impact on the surrounding environment (Tang and Maggi 2021; Riedo et al. 2022; Bošković et al. 2020). Studies looking at top soils from across Europe have revealed the prevalence of residues from a minimum of five pesticides in over half of the analyzed samples (Silva et al. 2019; Hvězdová et al. 2018; Riedo et al. 2021). These results indicate the widespread occurrence of multi-pesticide exposure and the considerable degree of anthropogenic pressure that could potentially be exerted on soil microbial communities.

Some microorganisms participate in pesticide degradation, thus reducing their concentrations in the environment (Mandelbaum, Wackett, and Allan 1993; Marrón-Montiel et al. 2006). It has been demonstrated, for instance, that bacterial genera such as *Achromobacter, Bacillus, Pseudomonas*, and *Rhodococcus* are particularly effective at breaking down pesticides. The transformation process is often enhanced by microbial co-metabolism, whereby byproducts from the metabolic transformations of one species can serve as substrates for others (Bælum, Jacobsen, and Holben 2010; Ermakova et al. 2010; Hou et al. 2011; Sandoval-Carrasco et al. 2013; Vargha, Takáts, and Márialigeti 2005; Zhang et al. 2011). However, the metabolic adaptability exhibited by these bacterial genera does not mean all microbes are protected from pesticide-induced stress. Due to the sheer variety of active substances present in pesticides, their effects on soil microbial communities remain unclear and context-dependent. While some studies suggest that herbicides such as atrazine do not induce persistent changes in microbial abundance, others report toxic effects, particularly from chlorinated herbicides such as diuron and linuron (Dennis et al. 2018; Prado and Airoldi 2002). Furthermore, the presence of pesticide residues in field sites has been shown to disrupt generic microbial metabolic pathways, reduce microbial biomass, and alter community composition (Ni et al. 2025; Riedo et al. 2021; Sim et al. 2022). It is thus clear that although some microorganisms can exploit pesticide residues as a resource, the extent to which pesticide accumulation and exposure alters microbial diversity and long-term community stability remains poorly understood (Rose et al. 2018; Banks et al. 2014).

The rate at which a global change agent is introduced into soils, also known as the rate of change, is an underexplored topic in ecology that potentially influences microbial community dynamics (Pinek et al. 2020). Although not abundantly, the relevance of global change agent exposure dynamics to microbial communities has already been established in a few studies. For instance, McTee et al. (2019) demonstrated that soil bacterial communities respond differently to abrupt versus gradual additions of copper, a heavy metal also commonly used in agriculture, with time playing a critical role in shaping the observed patterns. Similarly, it was shown that an abrupt increases in atmospheric CO_2_ can lead to an overestimation of community responses in a model plant-soil system, particularly among mycorrhizal fungi (Klironomos et al. 2005). While these studies provide valuable insights, they did not assess microbial community-level responses to pesticide exposure (Pinek et al. 2020). In real-world scenarios, pesticide accumulation frequently occurs in a gradual manner. Yet, most pesticide exposure studies focus on abrupt applications, overlooking these exposure patterns that more closely reflect environmental conditions (Golubeva et al. 2020; Li et al. 2021). Abrupt applications of pesticides have been demonstrated to potentially overwhelm microbial systems by introducing elevated concentrations of bioavailable compounds, thereby leading to oversaturation of the soil’s sorption sites (Meidl et al. 2024). In contrast, the application of pesticides on a gradual basis might allow microbial communities time to acclimate, although idiosyncratic responses have been observed (Golubeva et al. 2020). Clearly, understanding the role of exposure dynamics of pesticides in shaping microbial communities is critical for sustainable soil management, especially as pesticide-driven shifts in microbial community composition might have long-lasting effects on soil function.

The present study investigates the influence of pesticide application rates, whether abrupt or gradual, on soil microbial communities, with a particular focus on fungi and bacteria. We addressed two hypotheses: i) Among pesticide classes, fungicides will demonstrate the most profound effects on soil microbial communities when compared to herbicides and insecticides. This is particularly evident for fungal communities specifically designed to suppress fungi (Prudnikova, Streltsova, and Volova 2021). ii) Based on earlier results from this experimental setup, abrupt pesticide applications will have a more pronounced impact on soil microbial communities than gradual applications due to reduced adaptation time and increased bioavailability of the pesticides (Meidl et al. 2024).

## Material and Methods

### Experimental setup

A microcosm setup was used to evaluate the impact of diverse pesticides and application rates on soil communities. Details regarding the experimental setup can be found in a previous paper examining soil process-level responses using the same experiment (Meidl et al. 2024). In brief, the top 10 cm of surface soil was collected from a local grassland at an experimental site of the Freie Universität Berlin (52°46′60.67 N, 13°30′26.98 E). The soil was sieved (2 mm), and 30 grams were placed into micro-boxes (Deinze, Belgium; filter cover, 80 mm diameter, 40 mm height) and adjusted to a water holding capacity of 60%. The experiment was conducted over 50 days in a climate chamber with a humidity level of 30% and a temperature of 20°C. The ten pesticides included in the setup were selected based on their high prevalence in pesticide screening studies carried out in the European Union and Switzerland (Pelosi et al. 2021; Riedo et al. 2021), encompassing two insecticides, four fungicides, and four herbicides. The applied pesticide concentrations were based on the maximum detected concentrations across all 280 field sites (Table S1). The analytical standards (Sigma-Aldrich, United States) were diluted in acetone to create the working solutions. Two distinct treatment regimens were tested: an abrupt application, whereby the entire pesticide load was applied in a single dose, and a gradual application, whereby the dose was administered at five separate time points. The treatments were replicated ten times each. To evaluate the potential impact of solvents, the final concentration of acetone was incorporated as an additional control treatment in addition to the regular control comprising solely water. The treatments and controls were applied to the soil surface using a 1 mL glass syringe with a nasal spray pump (Teleflex Medical Europe Ltd., Ireland). The gradual treatments were applied at 10-day intervals, while the abrupt treatment was administered in a single dose on day 20. Following 50 days, soil samples were taken and stored in 2 mL Eppendorf tubes at -80°C until further analysis.

### Molecular analysis (DNA extraction, PCR amplification, and Illumina sequencing library preparation)

The DNA was extracted from 250 mg of lyophilized soil using the DNeasy Power Soil Pro Kit (Qiagen, GmbH, Germany), following the manufacturer’s instructions. The fungal sequences were amplified using the fITS7 and ITS4 primers, which, when used in combination, yield amplicons that span the fungal ITS2 region (Ihrmark et al. 2012). Bacterial sequences were amplified from soil DNA extracts using the 515F-Y and 806R primers (Caporaso et al. 2011). The polymerase chain reaction (PCR) amplification and sequencing library preparation steps were conducted on an automated workstation (Biomek i7 hybrid, Beckman Coulter, United States) at the Berlin Center for Genomics in Biodiversity Research (BeGenDiv, Berlin, Germany).

For the 16S target PCR, the reaction was carried out using Q5 High-Fidelity DNA polymerase (New England Biolabs, United States) in a 25-µl volume containing template DNA (10ng), 1.25 µl of both the 10 µM forward primer and 10 µM reverse primer, and 12.50 µl Q5 High-Fidelity 2X Master Mix. The target PCR comprised an initial denaturation step for 30 seconds at 98 °C, 30 cycles of a 10-second denaturation step at 98 °C, a 30-second annealing step at 57 °C, and an elongation step at 72 °C for 20 seconds. The final elongation step was conducted at 72°C for 120 seconds, after which the temperature was maintained at 4°C. The same reagents and concentrations were employed for the indexing PCR as were used for the previous reaction. The PCR program was set to 30 seconds at 98 °C for initiation, followed by eight PCR cycles (98 °C for 10 seconds, 67 °C for 30 seconds, 72 °C for 30 seconds) and a final elongation step at 72 °C for 120 seconds.

For ITS, the target PCR was conducted using Kapa HiFi polymerase (Roche, Switzerland). The reaction volume of 25 µl contained template DNA (20ng), 0.75 µl of 10 mM dNTPs (200 µM), 5 µl of Kapa HiFi buffer, 0.75 µl of both the 10 µM forward primer and 10 µM reverse primer, 0.5 µl of Kapa HiFi polymerase (0.02 U/µl), and nuclease-free water. The target PCR involved an initial denaturation step for 180 seconds at 98 °C, followed by 30 cycles of a 20-second denaturation step at 98 °C, a 20-second annealing step at 58 °C, and an elongation step at 72 °C for 30 seconds. The final elongation step lasted 300 seconds at 72 °C, after which the temperature was maintained at 4 °C. The indexing PCR was conducted using 10 µl purified DNA, 0.5 µl of 10 mM dNTPs (200 µM), 1.25 µl of both forward and reverse primers, 5 µl Kapa HiFi buffer, 0.5 µl Kapa HiFi polymerase (0.02 U/µl), and nuclease-free water in a 25 µl reaction volume. The PCR program was set to 180 seconds at 98 °C for initiation, followed by eight PCR cycles (98 °C for 20 seconds, 52 °C for 20 seconds, 72 °C for 30 seconds) and a final elongation step at 72 °C for 300 seconds.

Following each PCR, products were purified using magnetic beads (CleanNGS, GCbiotech, the Netherlands), and to ascertain the quality and efficacy of the indexing PCR, a randomized comparison was conducted between a target and index sample, using a Tape station (Agilent Technologies, United States). The products of the second PCR were then pooled for sequencing on a MiSeq System (Illumina, United States) with the V3 chemistry (600 cycles) and paired-end dual indexing. Following the sequencing process, the reads were demultiplexed and assigned to fastq-files by their indexes, utilizing the MiSeq software (Illumina, United States).

### Bioinformatics

The forward and reverse reads were processed separately using Dada2 v. 1.30.0 (Callahan et al. 2016), ShortRead v. 1.60.0 (Morgan et al. 2009), and Biostrings v. 2.70.1 (Pagès et al. 2021) to identify amplicon sequence variants (ASVs). The approach followed the Dada2 tutorial for ITS sequences (v1.8, https://benjjneb.github.io/dada2/ITS_workflow.html) and 16S sequences (v1.16, https://benjjneb.github.io/dada2/tutorial.html). First, primer sequences were removed from each read with Cutadapt v. 4.0 (Martin 2011). Reads were filtered to exclude any sequences under 50 base pairs (bp) in length and truncated where base quality scores dropped below a threshold of 10. Only reads with an expected error rate of 2 or lower were retained. Next, a machine-learning algorithm in Dada2 modeled the error frequency patterns across the filtered reads, and de-replication was used to identify unique sequence variants. The Dada2 sample inference algorithm then processed the reads based on learned error patterns, defining ASVs. Forward and reverse reads were merged, and any chimeras were removed. Finally, the raw reads were mapped back to the ASVs to generate an ASV abundance table. Taxonomic identity was assigned using the naive Bayesian classifier method of the DADA2 package against the UNITE database 9.0 (Abarenkov et al. 2024) for fungi and the Silva reference database v123 (Callahan 2018) for bacteria.

### Statistical analysis

All analyses were conducted using the R statistical computing environment, version 4.3.0. The data were organized and selected using the phyloseq package v1.46.0 (McMurdie and Holmes 2013), DESeq2 v1.42.1 (Love, Huber, and Anders 2014), and dplyr v1.14 (Wickham et al. 2023). All figures were generated using functions from the packages ggplot2 v3.5.1 (Wickham, Chang, and Wickham 2016).

The raw amplicon sequence variant (ASV) data were imported and subjected to preliminary processing, during which low-abundance ASVs and taxa not identified at the phylum level were excluded. The minimum sample depth was employed as the rarefaction threshold, with ASVs rarefied to this depth using the rarefy_even_depth function in phyloseq, with a fixed random seed (1782) to ensure reproducibility.

To evaluate microbial diversity within samples, alpha diversity metrics, including observed richness and inverse Simpson’s dominance, were computed. To estimate the mean and 95% confidence intervals (CI) of the response metrics for each treatment, a custom bootstrap resampling procedure was employed, as described by Rillig et al. (2019). For each treatment group, 1,000 bootstrap samples were generated through the resampling with replacement of the observed values. In each iteration of the resampling process, a mean was calculated, and the resulting bootstrap distribution was employed to derive the 2.5%, 50%, and 97.5% confidence intervals. Subsequently, a comparison was conducted between each treatment group and the control by subtracting the control means from the treatment means across the bootstrapped samples to estimate the effect size distribution’s 2.5%, 50%, and 97.5% percentiles. Furthermore, a significance level (p-value) was calculated with a Bonferroni correction applied to adjust for multiple comparisons. A density ridge plot was constructed to visually depict the raw data distributions, along with their bootstrapped confidence intervals.

The ASV data were subjected to a transformation process to facilitate the subsequent beta diversity analyses. A binary transformation (presence-absence) was applied to reduce the influence of dominant ASVs, thereby facilitating a robust beta diversity analysis. A Jaccard dissimilarity matrix was then calculated for the transformed ASV data. A Principal Coordinates Analysis (PCoA) was conducted using the Jaccard distance matrix (command vegdist() in package vegan), and the ordination scores were extracted for the first two axes. The PCoA results were visualized with a scatter plot, with ellipses representing the 95% confidence interval around each treatment group (control, gradual, abrupt). A PERMANOVA was performed to assess the differences between treatments and to determine whether these differences were due to differences in dispersion rather than true compositional differences, a dispersion test was performed using betadisper.

For the DESeq2 analysis, the data were filtered to exclude features with low counts (i.e., less than one count in over half of the samples) to reduce noise and focus on meaningful abundance patterns. Subsequently, the sample data were normalized using the internal methods of DESeq2 to account for discrepancies in sequencing depth. To identify significant differences in microbial abundance between treated and control samples, pairwise contrasts were performed. The resulting p-values were adjusted using the Benjamini-Hochberg procedure, and genera with adjusted p-values below 0.1 were considered significant. Heatmaps with the log2 fold change, relative abundance, and p-values were then created.

## Results

### Composition of the bacterial and fungal microbiota

A total of 4,383 ASVs originating from the 16S marker were identified. Following the rarefaction process, 442 of these variants were excluded. Despite this loss, the rarefaction curves indicated that the number of ASVs per sample approached saturation for the majority of samples at the selected threshold. Of the 3,941 16S-ASVs that remained, 3,680 were classified as bacterial. The most prevalent bacterial phyla were *Actinobacteria* (32.3%), *Proteobacteria* (19.4%), *Chloroflexi* (9.12%), *Firmicutes* (8.5%), *Thaumarchaeota* (7.9%), *Acidobacteria* (5.7%), *Planctomycetes* (4.4%), *Gemmatimonadetes* (3.7%), *Cyanobacteria* (2.7%), and *Verrucomicrobia* (2.6%). For ITS, the bioinformatic process defined a total of 2,091 ITS2-ASVs. Rarefaction resulted in the exclusion of 226 variants. All of the remaining 1,865 IT2S-ASVs were confirmed to be of fungal origin. Fungal ASVs were predominantly distributed across five major phyla and were *Ascomycota* (58.4%), *Basidiomycota* (15.7%), *Mortierellomycota* (12.4%), *Chytridiomycota* (4.4%), and *Mucoromycota* (2.7%).

### Effect of pesticide application on total bacterial and fungal diversity

Analysis based on the inverse Simpson’s dominance index (iSD) revealed that the diversity of bacterial communities was not significantly different from the control treatment (Figure 1, A). The mean effect size difference between the application of the treatments (gradual vs. abrupt) and the difference of richness estimates of all treatment groups in relation to the control were also not significant (Figure S1, A and Figure S2, A). Consequently, the remainder of this study will focus exclusively on the response of fungal communities.

**Figure 1:**
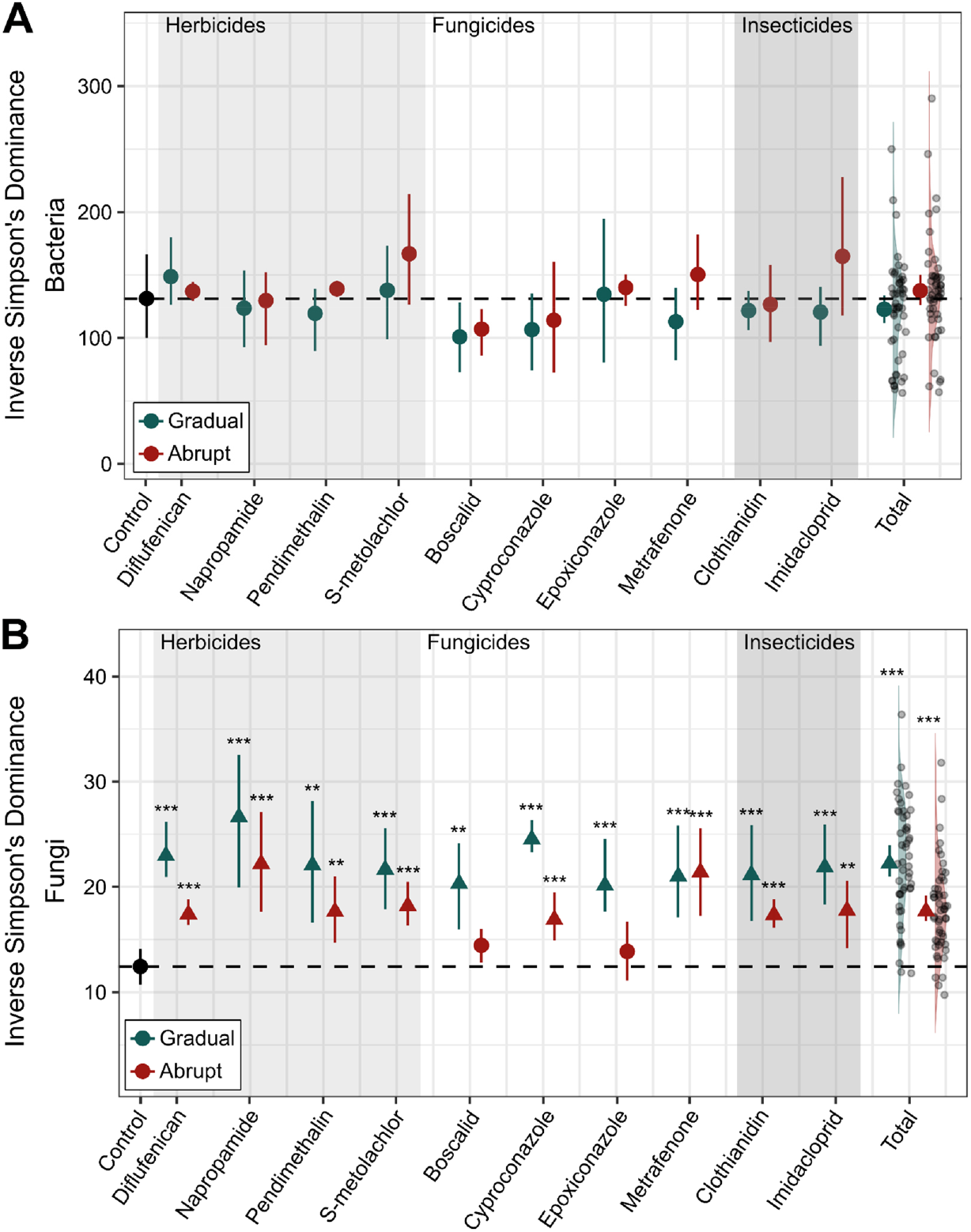
Individual effects of different pesticides (ordered by herbicides, fungicides, and insecticides) applied gradually (blue) or abruptly (red) on the inverse Simpson’s dominance index. The circles and triangles, along with the error bars, represent the mean effect size and 95% confidence intervals (CIs), respectively. The circles represent neutral effects, while the triangles (with an arrow pointing up or down) represent significant positive or negative effects. The asterisks mark the degree of significance (*: P-value < 0.05, **: P-value < 0.01, ***: P-Value < 0.001). For bacteria (A), no significant change in Simpson’s dominance index was observed for any of the ten treatments. For fungi (B), a significant increase in Simpson’s dominance index was observed for all treatments except the abrupt application of boscalid and epoxiconazole. A higher Inverse Simpson’s Dominance index reflects an increase in diversity and a more even distribution, reducing the dominance of few taxa.

The iSD of fungal communities significantly increased in almost all treatments, indicating an increase in the evenness of fungal communities (Figure 1, B). Exceptions to this pattern were observed for boscalid and epoxiconazole, where no significant changes occurred in the abrupt treatment compared to the control. Although, generally, pesticide application regimes caused significant increases in iSD in relation to the control group, differences in mean effect size between the application regimes (gradual vs. abrupt) could only be observed for diflufenican, cyproconazole, and epoxiconazole (Figure S2). Herbicides (diflufenican, napropamide, pendimethalin, and S-metolachlor) led to a significant increase in iSD compared to the control. The gradual application of herbicides consistently resulted in lower dominance than abrupt application. Similarly, gradual exposure to fungicides (boscalid, cyproconazole, epoxiconazole, and metrafenone) resulted in significantly higher evenness than abrupt exposure, with the notable exception of metrafenone. The broad-spectrum triazole fungicides cyproconazole and epoxiconazole showed particularly pronounced effects with gradual applications. Insecticides (clothianidin and imidacloprid) elicited mixed effects on iSD, with both gradual and abrupt applications increasing evenness relative to the control. Also, within this group of pesticides, gradual applications also consistently resulted in higher evenness than abrupt applications. The effects of pesticide treatments in general (summarized as total in Figure 1) showed increased iSD for both delivery methods.

Total fungal richness increments were observed only among four of the ten pesticides in the gradual treatment group, while only metrafenone showed a significant richness increase in the abrupt treatment group. Gradual delivery elicited significant richness increases across pesticide classes in relation to control (Figure S2, B). No notable patterns were observed regarding the relationship of fungal and bacterial communities with pesticide properties, such as DT_50_ or K_fOC_, which are responsible for the persistence of applied concentrations or the availability of substances (Table S2; data not shown).

### The impact of the rate of change on community diversity

Principal coordinate analysis (PCoA) revealed a clear clustering of treatment methods, indicating strong compositional dissimilarity between these two groups of communities and the control (Figure 2). For herbicides, fungal communities under both gradual and abrupt treatments are distinctly separated from the control in all cases (P-values < 0.05), suggesting shifts in community composition (Table S3). For diflufenican, napropamide, and S-metolachlor, the abrupt applications overlap with the gradual applications (P-values of 0.122, 0.149, and 0.055, respectively), indicating that there is no difference in the impact on fungal composition depending on the application method. With the exception of the abrupt epoxiconazole application (P-value = 0.161), fungicide treatments also induced clear shifts in fungal community structure, with gradual and abrupt applications forming distinct clusters separate from the control (P-values < 0.05). The broad-spectrum fungicide cyproconazole was the only fungicide treatment that elicited a distinct difference between the two treatment methods (P-value = 0.026). For the other fungicides, fungal communities under gradual applications overlap with those under abrupt applications (P-values > 0.05), indicating similar effects on the composition regardless of the application method. Both insecticides also induced shifts in the fungal community composition, leading to a clear separation of the two treatments from the control (P-values < 0.05). For clothianidin, the gradual and abrupt treatment clusters are not different from each other (P-value = 0.09), suggesting a more consistent mode of action across application methods.

**Figure 2:**
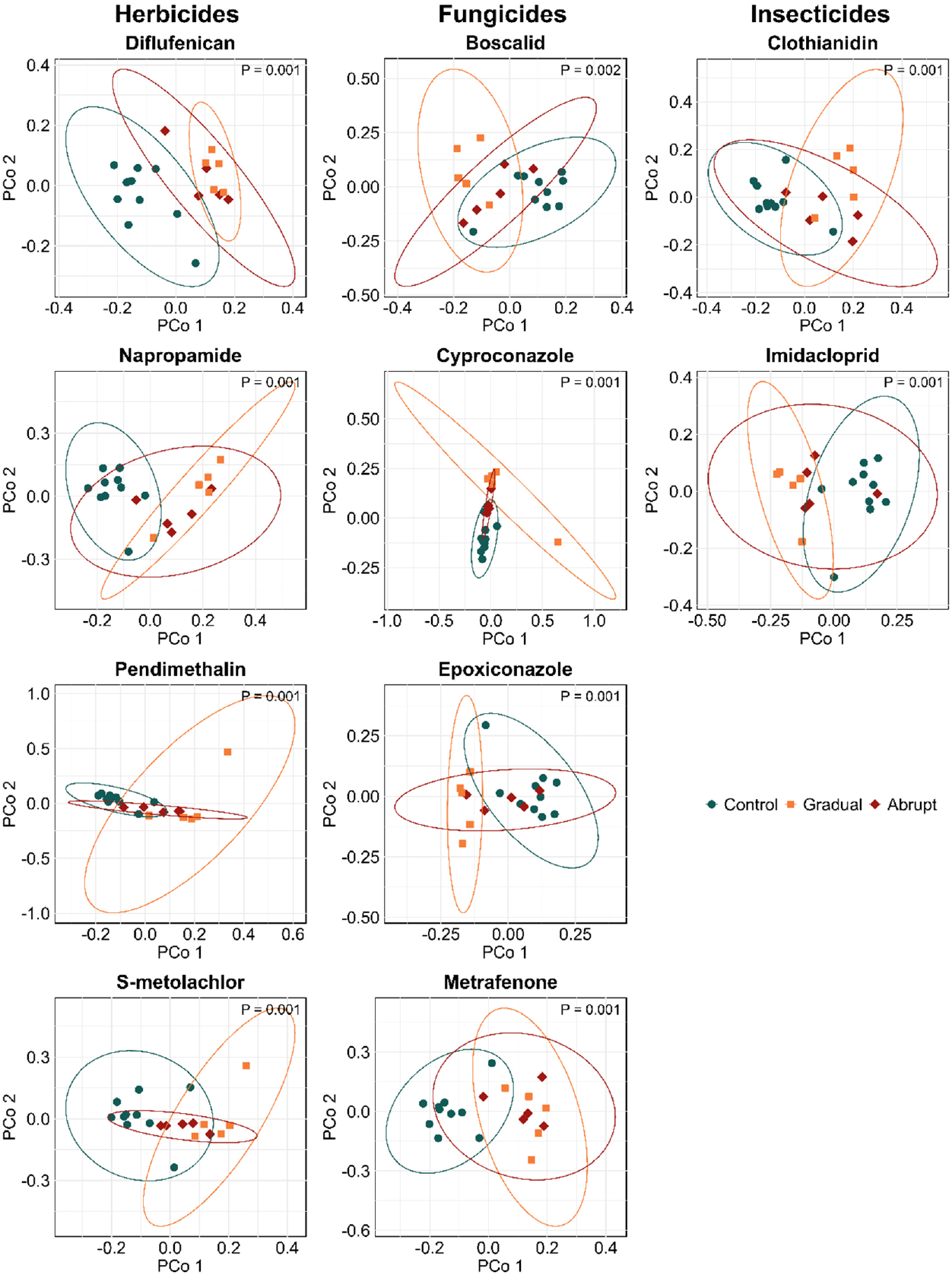
Principal coordinates analysis (PCoA) plot of the fungal species abundance based on the Jaccard distance of the control (blue dots), gradual application (orange squares), and abrupt application (red rhombuses). Each point represents a sample, and the distance between them indicates the compositional similarity. Ellipses correspond to clusters in each group with 95% confidence intervals. The P-value displayed on the plot represents the result of the overall PERMANOVA, indicating the significance of compositional differences between groups. A significant P-value (P < 0.05) suggests that treatment groups show distinct fungal community compositions.

### Influence of the pesticides on the abundance of individual genera

Differential abundance testing (expressed as Log2 fold changes) generally revealed that the most abundant fungal genera decreased in abundance when exposed to pesticides, while rare genera increased (Figure 3). Gradual herbicide treatments consistently resulted in larger positive or negative log2 fold changes than abrupt applications. For instance, the gradual application of pendimethalin significantly increased the abundance of genera like *Cladosporium* (+3.11, P-value < 0.001) and *Alternaria* (+2.92, P-value < 0.001). Abrupt treatments produced smaller shifts in relative genera abundance. For example, under abrupt napropamide application, genera such as *Hymenoscyphus* experienced a decrease in abundance (−5.54, P-value < 0.001), but the overall impact on relative abundance among other genera was less severe compared to gradual napropamide application. Fungicides, again, particularly broad-spectrum triazoles, had a pronounced effect on the relative abundance of fungal genera. For instance, under gradual cyproconazole treatment, *Tausonia* and *Enterocarpus* exhibited large relative abundance increases (+3.78 and +2.27, respectively, both P-value < 0.001). The supplementary heat map of relative abundances highlights the dominance of these genera (Figure S3). In contrast, abrupt fungicide treatments induced more transient disruptions, with smaller relative abundance shifts and fewer genera affected. Metrafenone, however, elicited consistent increases regardless of the application methods. Insecticides caused the least pronounced shifts overall, with fewer genera showing significant changes in abundance. Gradual insecticide applications resulted in modest shifts, with few genera responding to their presence. For example, under gradual clothianidin treatment, relative abundances across genera remained relatively stable. Abrupt insecticide application showed similarly limited effects on relative genera abundance.

**Figure 3:**
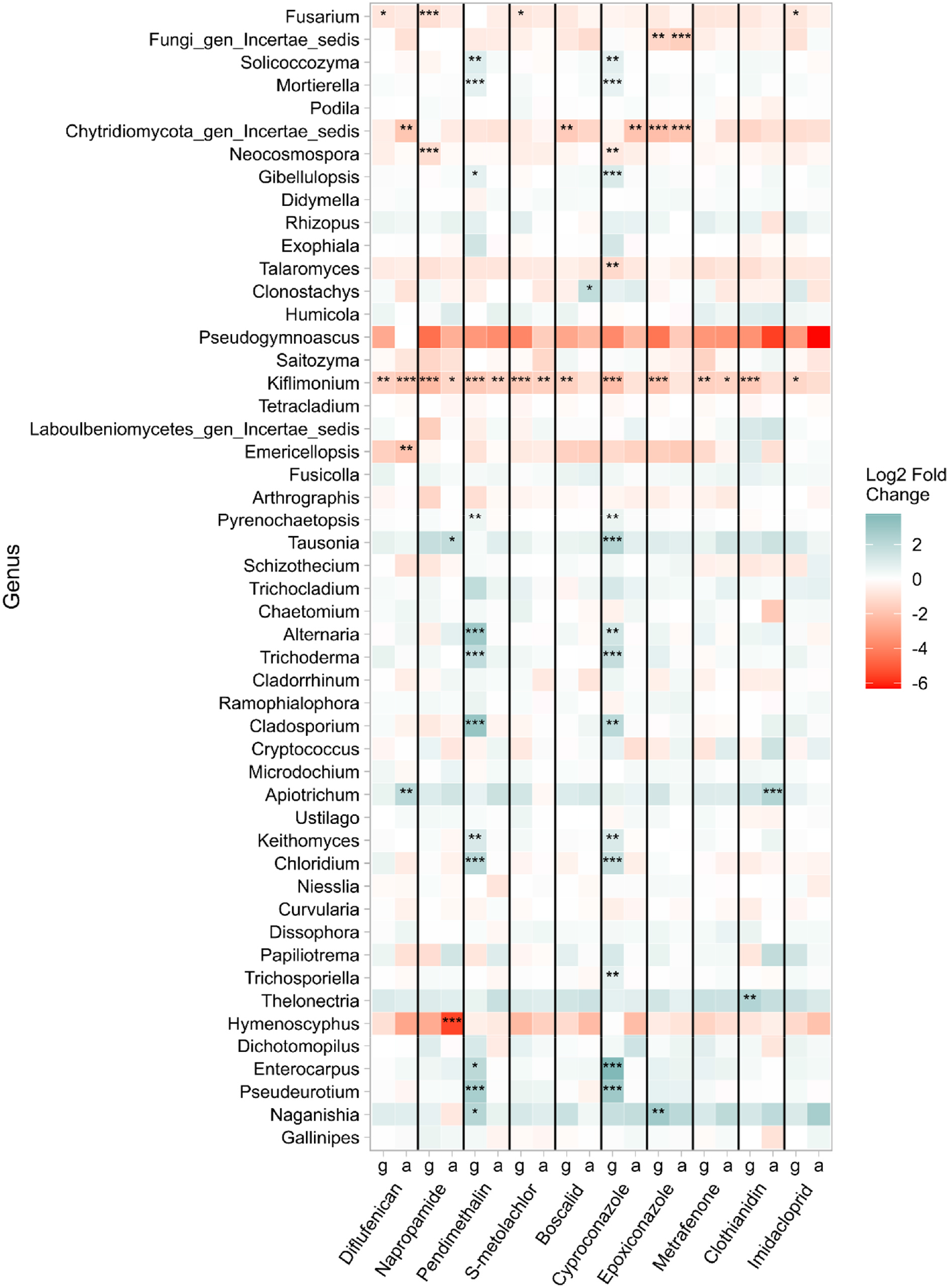
Heatmap showing the change in log2fold change of fungal genera as a function of pesticide application (g = gradual; a = abrupt) for the 50 most abundant genera, sorted by highest to lowest abundance. Red marks a decrease in abundance compared to the control, and blue marks an increase. The asterisks denote the degree of significance (P-value < 0.05, **: P-value < 0.01, ***: P-value < 0.001) in the change of abundance compared to the control.

## Discussion

Understanding how pesticide applications influence microbial communities is crucial for assessing their long-term ecological impacts. Yet, the overall relationship between microbial communities and pesticide exposure remains complex and poorly understood. Our study revealed that fungal communities were more responsive to pesticide applications than bacterial communities, which remained largely unaffected. Moreover, our findings provided mixed support for our hypotheses: Although fungicides had strong effects on fungal communities as expected, herbicides also induced considerable changes, suggesting broader non-target effects across pesticide classes. Additionally, gradual applications led to more pronounced changes in both alpha and beta diversity than abrupt applications, emphasizing the importance of exposure dynamics in shaping microbial responses.

While we first hypothesized that fungicides would have the most pronounced effects on soil microbial communities, our results indicate that pesticide effects in general were largely restricted to fungi, with bacterial communities showing little to no response. This suggests that fungi are generally more sensitive to pesticide exposure, aligning with previous research (Astaykina et al. 2020; Streletskii et al. 2022). The lack of effects on bacterial communities might be due to the relatively short duration of the experiment, as bacterial responses are often delayed and may only become apparent after longer exposure periods (Santísima-Trinidad et al. 2018; McTee et al. 2019). This lag could be due to differences in how fungi and bacteria interact with pesticides and their degradation products. Fungi are often the primary initiators of pesticide breakdown through extracellular enzymes, generating transformation products that may then be available to bacteria (Kumar et al. 2021; Katsoula et al. 2020). In other words, bacteria may benefit or suffer indirectly from pesticide exposure, with community-level shifts being detectable only once the resulting transformation products become available as substrates (Howell et al. 2014). Moreover, soil bacterial communities are known for their high diversity and metabolic plasticity, which may buffer short-term disturbances and mask compositional changes (Griffiths and Philippot 2013). Another possibility is that bacterial responses occur more subtly at the functional level or involve the proliferation of specific pesticide-degrading taxa without significantly altering overall community structure (Lo 2010). All these factors may explain the apparent insensitivity of bacterial communities to pesticide exposure over the timeframe of our study, highlighting the need for longer-term and functionally focused assessments when studying bacterial responses.

For fungal communities, our results partially support the expectation that fungicides would have the strongest effects, but herbicides also induced substantial changes in fungal diversity and composition. These observations suggest that pesticide impact cannot be solely attributed to differences in their mode of action (Prudnikova, Streltsova, and Volova 2021). For instance, the broad-spectrum fungicide cyproconazole led to pronounced shifts in fungal communities, consistent with its known inhibitory effects on major fungal taxa (Vasilchenko et al. 2023). However, herbicides such as napropamide and pendimethalin elicited equally strong responses, including increased fungal evenness and changes in the relative abundance of key taxa. These findings align with prior studies showing that some herbicides can selectively stimulate fungal taxa by acting as a substrate source for fungi capable of their degradation (Crouzet et al. 2016). Since no plants were included in our system, the observed herbicide effects must have been direct rather than indirectly mediated through plant-microbe interactions, further indicating that certain herbicides may exert selective pressures on fungal communities independent of their primary targets (Thiour-Mauprivez et al. 2019). This broader-than-expected consequence highlights the need to reconsider the assumption that non-fungicidal pesticides have insignificant effects on fungal communities.

Our second hypothesis predicted that abrupt pesticide applications would cause a more pronounced effect on microbial communities than gradual applications. However, contrary to this expectation, our results show that gradual pesticide application altered alpha diversity by increasing fungal evenness and beta diversity by shifting fungal species composition, with more distinct effects than abrupt application. This pattern likely stems from the ability of certain rare fungi to exploit selective conditions over time and thus adapt in the face of prolonged exposure. This argument becomes evident when looking at the effects of application rate at the genus level. Once again, gradual pesticide exposure resulted in more pronounced changes in both dominant and rare fungal genera, whereas abrupt applications resulted in smaller shifts in relative abundance. These results indicate that rare fungi may be occupying the ecological niches left by dominant fungi whose abundance has declined, resulting in reduced competition (Senabio et al. 2024; Roman et al. 2021). This also supports the notion that prolonged exposure creates a sustained selective pressure that enables certain taxa to thrive while suppressing the population growth of hyper-abundant genera. In our study, genera such as *Solicoccozyma* and *Trichoderma* increased in relative abundance under gradual treatment, suggesting a potential role in coping with chemical stress. Similar increases have been reported in previous studies, highlighting the potential role of these genera in pesticide biodegradation and resilience to xenobiotic compounds (Stosiek et al. 2019; Escudero-Leyva et al. 2022). The fact that only abundant taxa appear to be negatively affected by pesticide exposure might also be due to their widespread distribution, which might increase their likelihood of encountering pesticides. By contrast, rare taxa being more sparsely distributed may encounter pesticides less often (Stanley and Preetha 2016). Alternatively, the increase in fungal richness observed under some gradual treatments suggests that low pesticide doses may even elicit hormetic effects, potentially stimulating sporulation and the proliferation of rare fungal variants (Li et al. 2024; Agathokleous, Calabrese, and Veresoglou 2024). This phenomenon may further contribute to the observed shifts in fungal community composition, as certain taxa exploit pesticide-induced changes in resource availability or competition dynamics (Walder et al. 2022).

The pronounced shifts in fungal diversity and composition observed under gradual pesticide exposure may have important ecological implications (Wang et al. 2024). Fungal community shifts could impact key ecosystem processes such as carbon turnover and nitrogen cycling, particularly if the emergent fungal taxa possess different enzymatic capabilities or metabolic pathways (Bardgett and van der Putten 2014; Wagg et al. 2014; Schimel 2016). In this regard, our results complement prior studies, as they indicate that shifts in fungal community attributes can indeed be correlated with strong shifts in certain soil processes. Meidl et al. (2024), analyzing data from the same experiment, reported that individual pesticide addition increased decomposition rates, as well as measures of soil aggregate stability. Both organic matter decomposition and soil aggregation are processes to which soil fungi are fundamental agents (Gessner et al. 2010; Rillig and Mummey 2006). Furthermore, even though we observed that the gradual application rate exerted stronger effects on fungal communities relative to abrupt application, our results appear to indicate that even modest fungal community shifts could lead to sizable changes in the soil processes to which these organisms are directly relevant. These results also align with previous research showing that repeated low-dose pesticide exposure can impose long-term selective pressures on microbial communities (Biache et al. 2017), reducing the window for recovery and increasing cumulative effects, thereby acting as sustained selective pressures that reshape fungal communities more profoundly (Zhang et al. 2016; Druille et al. 2016). However, despite our results suggesting that shifts in soil functions elicited by individual pesticide exposure can be coupled with shifts in fungal community attributes, we cannot conclusively draw a causal link between these two phenomena.

A limitation of our study is that it does not allow us to understand the long-term trends of pesticide exposure on soil microbial communities. While we have shown that pesticide applications may cause short-term disturbances to fungal communities, it remains unclear whether fungal communities, or the functions these govern, eventually return to their original state or remain altered after pesticide exposure ceases. If previously dominant taxa fail to recover, the rate and stability of key ecosystem processes may be permanently affected, as implied by the insurance hypothesis, which suggests that higher biodiversity can stabilize ecosystem functioning in the face of environmental change by ensuring that some species compensate when others decline (Naeem and Li 1997; Yachi and Loreau 1999; Griffiths and Philippot 2013). Over time, such selective pressures on microbial communities may permanently reshape soil functions, potentially leading to fundamental changes in nutrient dynamics and overall ecosystem stability (Groffman et al. 2006). From an agricultural perspective, our findings highlight the need to consider both application rate and frequency when assessing pesticide impacts. While most pesticide assessments are based on single-dose exposures, our results suggest that repeated low-dose exposures may have more persistent and selective effects on microbial communities (Riedo, Rillig, and Walder 2025). Incorporating gradual exposure scenarios into longer-term ecotoxicological studies could provide a more accurate representation of real-world pesticide dynamics and their long-term effects on soil microbiota. To better capture these dynamics, future research should include more than two levels of pesticide application intensity (Pinek et al. 2020). Doing so would enable a more nuanced analysis of both the rate of change and the mode of exposure, bridging the current gap between dose-response and temporally explicit experimental designs.

## Conclusion

This study emphasizes the influence of pesticide application methods on soil microbial communities. Our findings confirm that fungal communities are more sensitive to pesticide exposure than bacterial communities, exhibiting a more pronounced shift in both alpha and beta diversity. Gradual applications promoted the proliferation of fungal taxa with lower abundances, particularly under certain fungicides and herbicides, while abrupt treatments caused less severe disruptions. These results suggest that the frequency of low-dosage pesticide exposure can drive profound changes in fungal community composition, potentially impacting soil health and ecosystem functions. Long-term studies are needed to capture potentially delayed bacterial responses and to assess how chronic, low-dose pesticide exposure affects the recovery of microbial communities. Understanding whether microbial communities can recover or remain permanently altered is essential for predicting long-term ecosystem consequences and informing sustainable soil management practices. Future research should also explore whether observed shifts in fungal diversity translate into functional changes, such as altered decomposition rates or nutrient cycling. Moving beyond taxonomic shifts to examine microbial functional traits could provide deeper mechanistic insights into how pesticides reshape soil ecosystems. An improved understanding of these dynamics is thus crucial for developing sustainable pesticide application strategies that minimize unintended ecological consequences.

## CRediT Authorship Contribution Statement

**Judith Riedo:** Conceptualization, Data curation, Formal analysis, Funding acquisition, Methodology, Visualization, Writing – original draft, Writing – review & editing. **Juan F. Dueñas**: Data curation, Formal analysis, Writing – review & editing. **Susan Mbedi:** Investigation, Writing – review & editing. **Sarah Sparmann:** Investigation, Writing – review & editing. **Matthias C. Rillig:** Conceptualization, Funding acquisition, Project administration, Resources, Writing – review & editing

## Data Availability

All sequences derived from this study are available at the European Nucleotide Archive (ENA) under project PRJEB88969. Raw reads are available with accessions ERR14895225-ERR14895545. Sample metadata are available with accessions ERS24236319-ERS24236478.

## References

[1] Abarenkov, Kessy, R Henrik Nilsson, Karl-Henrik Larsson, Andy F S Taylor, Tom W May, Tobias Guldberg Frøslev, Julia Pawlowska, et al. 2024. ‘The UNITE Database for Molecular Identification and Taxonomic Communication of Fungi and Other Eukaryotes: Sequences, Taxa and Classifications Reconsidered’. Nucleic Acids Research 52 (D1): D791–97. 10.1093/nar/gkad1039.

[2] Agathokleous, Evgenios, Edward J. Calabrese, and Stavros D. Veresoglou. 2024. ‘The Microbiome Orchestrates Contaminant Low-Dose Phytostimulation’. Trends in Plant Science, December. 10.1016/j.tplants.2024.11.019.

[3] Astaykina, A. A., R. A. Streletskii, M. N. Maslov, A. A. Belov, V. S. Gorbatov, and A. L. Stepanov. 2020. ‘The Impact of Pesticides on the Microbial Community of Agrosoddy-Podzolic Soil’. Eurasian Soil Science 53 (5): 696–706. 10.1134/S1064229320050038.

[4] Bælum, Jacob, Carsten S. Jacobsen, and William E. Holben. 2010. ‘Comparison of 16S rRNA Gene Phylogeny and Functional tfdA Gene Distribution in Thirty-One Different 2,4-Dichlorophenoxyacetic Acid and 4-Chloro-2-Methylphenoxyacetic Acid Degraders’. Systematic and Applied Microbiology 33 (2): 67–70. 10.1016/j.syapm.2010.01.001.

[5] Banks, M. L., A. C. Kennedy, R. J. Kremer, and F. Eivazi. 2014. ‘Soil Microbial Community Response to Surfactants and Herbicides in Two Soils’. Applied Soil Ecology 74 (February):12–20. 10.1016/j.apsoil.2013.08.018.

[6] Bardgett, Richard D., and Wim H. van der Putten. 2014. ‘Belowground Biodiversity and Ecosystem Functioning’. Nature 515 (7528): 505–11. 10.1038/nature13855.

[7] Biache, Coralie, Salma Ouali, Aurélie Cébron, Catherine Lorgeoux, Stéfan Colombano, and Pierre Faure. 2017. ‘Bioremediation of PAH-Contamined Soils: Consequences on Formation and Degradation of Polar-Polycyclic Aromatic Compounds and Microbial Community Abundance’. Journal of Hazardous Materials 329 (May):1–10. 10.1016/j.jhazmat.2017.01.026.

[8] Bošković, Nikola, Kerstin Brandstätter-Scherr, Petr Sedláček, Zuzana Bílková, Lucie Bielská, and Jakub Hofman. 2020. ‘Adsorption of Epoxiconazole and Tebuconazole in Twenty Different Agricultural Soils in Relation to Their Properties’. Chemosphere 261 (December):127637. 10.1016/j.chemosphere.2020.127637.

[9] Callahan, Benjamin. 2018. ‘Silva Taxonomic Training Data Formatted for DADA2 (Silva Version 132)’. Zenodo. 10.5281/zenodo.1172783.

[10] Callahan, Benjamin J., Paul J. McMurdie, Michael J. Rosen, Andrew W. Han, Amy Jo A. Johnson, and Susan P. Holmes. 2016. ‘DADA2: High-Resolution Sample Inference from Illumina Amplicon Data’. Nature Methods 13 (7): 581–83. 10.1038/nmeth.3869.

[11] Caporaso, J. Gregory, Christian L. Lauber, William A. Walters, Donna Berg-Lyons, Catherine A. Lozupone, Peter J. Turnbaugh, Noah Fierer, and Rob Knight. 2011. ‘Global Patterns of 16S rRNA Diversity at a Depth of Millions of Sequences per Sample’. Proceedings of the National Academy of Sciences 108 (upplement_1): 4516–22. 10.1073/pnas.1000080107.

[12] Crouzet, Olivier, Franck Poly, Frédérique Bonnemoy, David Bru, Isabelle Batisson, Jacques Bohatier, Laurent Philippot, and Clarisse Mallet. 2016. ‘Functional and Structural Responses of Soil N-Cycling Microbial Communities to the Herbicide Mesotrione: A Dose-Effect Microcosm Approach’. Environmental Science and Pollution Research 23 (5): 4207–17. 10.1007/s11356-015-4797-8.

[13] Dennis, Paul G., Tegan Kukulies, Christian Forstner, Thomas G. Orton, and Anthony B. Pattison. 2018. ‘The Effects of Glyphosate, Glufosinate, Paraquat and Paraquat-Diquat on Soil Microbial Activity and Bacterial, Archaeal and Nematode Diversity’. Scientific Reports 8 (1): 2119. 10.1038/s41598-018-20589-6.

[14] Druille, M., P.A. García-Parisi, R. A. Golluscio, F. P. Cavagnaro, and M. Omacini. 2016. ‘Repeated Annual Glyphosate Applications May Impair Beneficial Soil Microorganisms in Temperate Grassland’. Agriculture, Ecosystems & Environment 230 (August):184–90. 10.1016/j.agee.2016.06.011.

[15] Ermakova, Inna T., Nina I. Kiseleva, Tatyana Shushkova, Mikhail Zharikov, Gennady A. Zharikov, and Alexey A. Leontievsky. 2010. ‘Bioremediation of Glyphosate-Contaminated Soils’. Applied Microbiology and Biotechnology 88 (2): 585–94. 10.1007/s00253-010-2775-0.

[16] Escudero-Leyva, Efraín, Pamela Alfaro-Vargas, Rodrigo Muñoz-Arrieta, Camila Charpentier-Alfaro, María del Milagro Granados-Montero, Katherine S. Valverde-Madrigal, Marta Pérez-Villanueva, et al. 2022. ‘Tolerance and Biological Removal of Fungicides by Trichoderma Species Isolated From the Endosphere of Wild Rubiaceae Plants’. Frontiers in Agronomy 3 (February). 10.3389/fagro.2021.772170.

[17] European Commission. 2020. ‘EU Pesticides Database’. 2020. https://food.ec.europa.eu/plants/pesticides/eu-pesticides-database_en.

[18] FAO. 2023. Pesticides Use and Trade 1990–2021. FAOSTAT Analytical Briefs, No. 70. Rome, Italy: FAO. 10.4060/cc6958ens.

[19] Fierer, Noah. 2017. ‘Embracing the Unknown: Disentangling the Complexities of the Soil Microbiome’. Nature Reviews Microbiology 15 (10): 579–90. 10.1038/nrmicro.2017.87.

[20] Gessner, Mark O., Christopher M. Swan, Christian K. Dang, Brendan G. McKie, Richard D. Bardgett, Diana H. Wall, and Stephan Hättenschwiler. 2010. ‘Diversity Meets Decomposition’. Trends in Ecology & Evolution 25 (6): 372–80. 10.1016/j.tree.2010.01.010.

[21] Golubeva, Polina, Masahiro Ryo, Ludo A. H. Muller, Max-Bernhard Ballhausen, Anika Lehmann, Moisés A. Sosa-Hernández, and Matthias C. Rillig. 2020. ‘Soil Saprobic Fungi Differ in Their Response to Gradually and Abruptly Delivered Copper’. Frontiers in Microbiology 11 (June):1195. 10.3389/fmicb.2020.01195.

[22] Griffiths, Bryan S., and Laurent Philippot. 2013. ‘Insights into the Resistance and Resilience of the Soil Microbial Community’. FEMS Microbiology Reviews 37 (2): 112–29. 10.1111/j.1574-6976.2012.00343.x.

[23] Groffman, Peter M., Jill S. Baron, Tamara Blett, Arthur J. Gold, Iris Goodman, Lance H. Gunderson, Barbara M. Levinson, et al. 2006. ‘Ecological Thresholds: The Key to Successful Environmental Management or an Important Concept with No Practical Application?’ Ecosystems 9 (1): 1–13. 10.1007/s10021-003-0142-z.

[24] Hou, Ying, Jian Tao, Wenjing Shen, Juan Liu, Jingquan Li, Yongfeng Li, Hui Cao, and Zhongli Cui. 2011. ‘Isolation of the Fenoxaprop-Ethyl (FE)-Degrading Bacterium Rhodococcus Sp. T1, and Cloning of FE Hydrolase Gene Feh’. FEMS Microbiology Letters 323 (2): 196–203. 10.1111/j.1574-6968.2011.02376.x.

[25] Howell, Christopher C., Sally Hilton, Kirk T. Semple, and Gary D. Bending. 2014. ‘Resistance and Resilience Responses of a Range of Soil Eukaryote and Bacterial Taxa to Fungicide Application’. Chemosphere 112 (October):194–202. 10.1016/j.chemosphere.2014.03.031.

[26] Hvězdová, Martina, Petra Kosubová, Monika Košíková, Kerstin E. Scherr, Zdeněk Šimek, Lukáš Brodský, Marek Šudoma, et al. 2018. ‘Currently and Recently Used Pesticides in Central European Arable Soils’. Science of The Total Environment 613–614 (February):361–70. 10.1016/j.scitotenv.2017.09.049.

[27] Ihrmark, Katarina, Inga T.M. Bödeker, Karelyn Cruz-Martinez, Hanna Friberg, Ariana Kubartova, Jessica Schenck, Ylva Strid, et al. 2012. ‘New Primers to Amplify the Fungal ITS2 Region – Evaluation by 454-Sequencing of Artificial and Natural Communities’. FEMS Microbiology Ecology 82 (3): 666–77. 10.1111/j.1574-6941.2012.01437.x.

[28] Katsoula, A, S Vasileiadis, M Sapountzi, and Dimitrios G Karpouzas. 2020. ‘The Response of Soil and Phyllosphere Microbial Communities to Repeated Application of the Fungicide Iprodione: Accelerated Biodegradation or Toxicity?’ FEMS Microbiology Ecology 96 (6): fiaa056. 10.1093/femsec/fiaa056.

[29] Klironomos, John N., Michael F. Allen, Matthias C. Rillig, Jeff Piotrowski, Shokouh Makvandi-Nejad, Benjamin E. Wolfe, and Jeff R. Powell. 2005. ‘Abrupt Rise in Atmospheric CO2 Overestimates Community Response in a Model Plant–Soil System’. Nature 433 (7026): 621–24. 10.1038/nature03268.

[30] Kumar, Manish, Ajar Nath Yadav, Raghvendra Saxena, Diby Paul, and Rajesh Singh Tomar. 2021. ‘Biodiversity of Pesticides Degrading Microbial Communities and Their Environmental Impact’. Biocatalysis and Agricultural Biotechnology 31 (January):101883. 10.1016/j.bcab.2020.101883.

[31] Lehmann, Anika, Weishuang Zheng, and Matthias C. Rillig. 2017. ‘Soil Biota Contributions to Soil Aggregation’. Nature Ecology & Evolution 1 (12): 1828–35. 10.1038/s41559-017-0344-y.

[32] Li, Erqin, Aleksandra Krsmanovic, Max-Bernhard Ballhausen, and Matthias C. Rillig. 2021. ‘Fungal Response to Abruptly or Gradually Delivered Antifungal Agent Amphotericin B Is Growth Stage Dependent’. Environmental Microbiology 23 (12): 7701–9. 10.1111/1462-2920.15797.

[33] Li, Yong, Kaiwei Zhang, Jian Chen, Leigang Zhang, Fayun Feng, Jinjin Cheng, Liya Ma, et al. 2024. ‘Rhizosphere Bacteria Help to Compensate for Pesticide-Induced Stress in Plants’. Environmental Science & Technology 58 (28): 12542–53. 10.1021/acs.est.4c04196.

[34] Lo, Chi-Chu. 2010. ‘Effect of Pesticides on Soil Microbial Community’. Journal of Environmental Science and Health, Part B 45 (5): 348–59. 10.1080/03601231003799804.

[35] Love, Michael I, Wolfgang Huber, and Simon Anders. 2014. ‘Moderated Estimation of Fold Change and Dispersion for RNA-Seq Data with DESeq2’. Genome Biology 15 (12): 550. 10.1186/s13059-014-0550-8.

[36] Mandelbaum, R. T., L. P. Wackett, and D. L. Allan. 1993. ‘Mineralization of the S-Triazine Ring of Atrazine by Stable Bacterial Mixed Cultures’. Applied and Environmental Microbiology 59 (6): 1695–1701. 10.1128/aem.59.6.1695-1701.1993.

[37] Marrón-Montiel, E., N. Ruiz-Ordaz, C. Rubio-Granados, C. Juárez-Ramírez, and C.J. Galíndez-Mayer. 2006. ‘2,4-D-Degrading Bacterial Consortium: Isolation, Kinetic Characterization in Batch and Continuous Culture and Application for Bioaugmenting an Activated Sludge Microbial Community’. Process Biochemistry 41 (7): 1521–28. 10.1016/j.procbio.2006.02.012.

[38] Martin, Marcel. 2011. ‘Cutadapt Removes Adapter Sequences from High-Throughput Sequencing Reads’. EMBnet.Journal 17 (1): 10–12. 10.14806/ej.17.1.200.

[39] McMurdie, Paul J., and Susan Holmes. 2013. ‘Phyloseq: An R Package for Reproducible Interactive Analysis and Graphics of Microbiome Census Data’. PLOS ONE 8 (4): e61217. 10.1371/journal.pone.0061217.

[40] McTee, Michael, Lorinda Bullington, Matthias C Rillig, and Philip W Ramsey. 2019. ‘Do Soil Bacterial Communities Respond Differently to Abrupt or Gradual Additions of Copper?’ FEMS Microbiology Ecology 95 (1): fiy212. 10.1093/femsec/fiy212.

[41] Meidl, Peter, Anika Lehmann, Mohan Bi, Carla Breitenreiter, Jasmina Benkrama, Erqin Li, Judith Riedo, and Matthias C. Rillig. 2024. ‘Combined Application of up to Ten Pesticides Decreases Key Soil Processes’. Environmental Science and Pollution Research 31 (8): 11995–4. 10.1007/s11356-024-31836-x.

[42] Morgan, Martin, Simon Anders, Michael Lawrence, Patrick Aboyoun, Hervé Pagès, and Robert Gentleman. 2009. ‘ShortRead: A Bioconductor Package for Input, Quality Assessment and Exploration of High-Throughput Sequence Data’. Bioinformatics 25 (19): 2607–8. 10.1093/bioinformatics/btp450.

[43] Naeem, Shahid, and Shibin Li. 1997. ‘Biodiversity Enhances Ecosystem Reliability’. Nature 390 (6659): 507–9. 10.1038/37348.

[44] Ni, Bang, Lu Xiao, D. Lin, Tian-Lun Zhang, Qi Zhang, Yanjie Liu, Quan Chen, et al. 2025. ‘Increasing Pesticide Diversity Impairs Soil Microbial Functions’. Proceedings of the National Academy of Sciences 122 (2): e2419917122. 10.1073/pnas.2419917122.

[45] Pagès, H, P Aboyoun, R Gentleman, and S DebRoy. 2021. ‘Biostrings: Efficient Manipulation of Biological Strings R Package Version 2.62. 0.(2021) https://Bioconductor.Org/Packages’. Biostrings.

[46] Pelosi, C., C. Bertrand, G. Daniele, M. Coeurdassier, P. Benoit, S. Nélieu, F. Lafay, et al. 2021. ‘Residues of Currently Used Pesticides in Soils and Earthworms: A Silent Threat?’ Agriculture, Ecosystems & Environment 305 (January):107167. 10.1016/j.agee.2020.107167.

[47] Pinek, Liliana, India Mansour, Milica Lakovic, Masahiro Ryo, and Matthias C. Rillig. 2020. ‘Rate of Environmental Change across Scales in Ecology’. Biological Reviews 95 (6): 1798–1811. 10.1111/brv.12639.

[48] Prado, Alexandre G. S., and Claudio Airoldi. 2002. ‘The Toxic Effect on Soil Microbial Activity Caused by the Free or Immobilized Pesticide Diuron’. Thermochimica Acta, 394 (1): 155–62. 10.1016/S0040-6031(02)00265-4.

[49] Prudnikova, Svetlana, Nadezhda Streltsova, and Tatiana Volova. 2021. ‘The Effect of the Pesticide Delivery Method on the Microbial Community of Field Soil’. Environmental Science and Pollution Research 28 (7): 8681–97. 10.1007/s11356-020-11228-7.

[50] Riedo, Judith, Chantal Herzog, Samiran Banerjee, Kathrin Fenner, Florian Walder, Marcel G.A. van der Heijden, and Thomas D. Bucheli. 2022. ‘Concerted Evaluation of Pesticides in Soils of Extensive Grassland Sites and Organic and Conventional Vegetable Fields Facilitates the Identification of Major Input Processes’. Environmental Science & Technology 56 (19): 13686–95. 10.1021/acs.est.2c02413.

[51] Riedo, Judith, Matthias C. Rillig, and Florian Walder. 2025. ‘Beyond Dosage: The Need for More Realistic Research Scenarios to Understand Pesticide Impacts on Agricultural Soils’. Journal of Agricultural and Food Chemistry, April. 10.1021/acs.jafc.4c12818.

[52] Riedo, Judith, Felix E. Wettstein, Andrea Rösch, Chantal Herzog, Samiran Banerjee, Lucie Büchi, Raphaël Charles, et al. 2021. ‘Widespread Occurrence of Pesticides in Organically Managed Agricultural Soils—the Ghost of a Conventional Agricultural Past?’ Environmental Science & Technology 55 (5): 2919–28. 10.1021/acs.est.0c06405.

[53] Rillig, Matthias C., and Daniel L. Mummey. 2006. ‘Mycorrhizas and Soil Structure’. New Phytologist 171 (1): 41–53. 10.1111/j.1469-8137.2006.01750.x.

[54] Rillig, Matthias C., Masahiro Ryo, Anika Lehmann, Carlos A. Aguilar-Trigueros, Sabine Buchert, Anja Wulf, Aiko Iwasaki, Julien Roy, and Gaowen Yang. 2019. ‘The Role of Multiple Global Change Factors in Driving Soil Functions and Microbial Biodiversity’. Science 366 (6467): 886–90. 10.1126/science.aay2832.

[55] Roman, Diana Larisa, Denisa Ioana Voiculescu, Madalina Filip, Vasile Ostafe, and Adriana Isvoran. 2021. ‘Effects of Triazole Fungicides on Soil Microbiota and on the Activities of Enzymes Found in Soil: A Review’. Agriculture 11 (9): 893. 10.3390/agriculture11090893.

[56] Rose, Michael T., Ee Ling Ng, Zhe (Han) Weng, Rachel Wood, Terry J. Rose, and Lukas Van Zwieten. 2018. ‘Minor Effects of Herbicides on Microbial Activity in Agricultural Soils Are Detected by N-Transformation but Not Enzyme Activity Assays’. European Journal of Soil Biology 87 (May):72– 79. 10.1016/j.ejsobi.2018.04.003.

[57] Sandoval-Carrasco, Carlos A., Deifilia Ahuatzi-Chacón, Juvencio Galíndez-Mayer, Nora Ruiz-Ordaz, Cleotilde Juárez-Ramírez, and Fernando Martínez-Jerónimo. 2013. ‘Biodegradation of a Mixture of the Herbicides Ametryn, and 2,4-Dichlorophenoxyacetic Acid (2,4-D) in a Compartmentalized Biofilm Reactor’. Bioresource Technology, Special Issue: IBS 2012 & Special Issue: IFIBiop, 145 (October):33–36. 10.1016/j.biortech.2013.02.068.

[58] Santísima-Trinidad, Ana Belén López, María del Mar Montiel-Rozas, Miguel Ángel Diéz-Rojo, Jose Antonio Pascual, and Margarita Ros. 2018. ‘Impact of Foliar Fungicides on Target and Non-Target Soil Microbial Communities in Cucumber Crops’. Ecotoxicology and Environmental Safety 166 (December):78–85. 10.1016/j.ecoenv.2018.09.074.

[59] Schimel, Joshua. 2016. ‘Microbial Ecology: Linking Omics to Biogeochemistry’. Nature Microbiology 1 (January):15028. 10.1038/nmicrobiol.2015.28.

[60] Senabio, Jaqueline Alves, Rafael Correia da Silva, Daniel Guariz Pinheiro, Leonardo Gomes de Vasconcelos, and Marcos Antônio Soares. 2024. ‘The Pesticides Carbofuran and Picloram Alter the Diversity and Abundance of Soil Microbial Communities’. PLOS ONE 19 (11): e0314492. 10.1371/journal.pone.0314492.

[61] Silva, Vera, Hans G. J. Mol, Paul Zomer, Marc Tienstra, Coen J. Ritsema, and Violette Geissen. 2019. ‘Pesticide Residues in European Agricultural Soils – A Hidden Reality Unfolded’. Science of The Total Environment 653 (February):1532–45. 10.1016/j.scitotenv.2018.10.441.

[62] Sim, Jowenna X. F., Casey L. Doolette, Sotirios Vasileiadis, Barbara Drigo, Ethan R. Wyrsch, Steven P. Djordjevic, Erica Donner, Dimitrios G. Karpouzas, and Enzo Lombi. 2022. ‘Pesticide Effects on Nitrogen Cycle Related Microbial Functions and Community Composition’. Science of The Total Environment 807 (February):150734. 10.1016/j.scitotenv.2021.150734.

[63] Stanley, Johnson, and Gnanadhas Preetha. 2016. ‘Pesticide Toxicity to Microorganisms: Exposure, Toxicity and Risk Assessment Methodologies’. In Pesticide Toxicity to Non-Target Organisms: Exposure, Toxicity and Risk Assessment Methodologies, edited by Johnson Stanley and Gnanadhas Preetha, 351–410. Dordrecht: Springer Netherlands. 10.1007/978-94-017-7752-0_6.

[64] Stosiek, Natalia, Agata Terebieniec, Adam Ząbek, Piotr Mlynarz, Hubert Cieslinski, and Magdalena Klimek-Ochab. 2019. ‘N-Phosphonomethylglycine Utilization by the Psychrotolerant Yeast Solicoccozyma Terricola M 3.1.4.’ Bioorganic Chemistry 93 (December):102866. 10.1016/j.bioorg.2019.03.040.

[65] Streletskii, Rostislav, Angelika Astaykina, George Krasnov, and Victor Gorbatov. 2022. ‘Changes in Bacterial and Fungal Community of Soil under Treatment of Pesticides’. Agronomy 12 (1): 124. 10.3390/agronomy12010124.

[66] Tang, Fiona H. M., and Federico Maggi. 2021. ‘Pesticide Mixtures in Soil: A Global Outlook’. Environmental Research Letters 16 (4): 044051. 10.1088/1748-9326/abe5d6.

[67] Thiour-Mauprivez, Clémence, Fabrice Martin-Laurent, Christophe Calvayrac, and Lise Barthelmebs. 2019. ‘Effects of Herbicide on Non-Target Microorganisms: Towards a New Class of Biomarkers?’ Science of The Total Environment 684 (September):314–25. 10.1016/j.scitotenv.2019.05.230.

[68] Vargha, Márta, Zoltán Takáts, and Károly Márialigeti. 2005. ‘Degradation of Atrazine in a Laboratory Scale Model System with Danube River Sediment’. Water Research 39 (8): 1560–68. 10.1016/j.watres.2004.10.013.

[69] Vasilchenko, Anastasia V., Darya V. Poshvina, Mikhail V. Semenov, Vyacheslav N. Timofeev, Alexandr V. Iashnikov, Artyom A. Stepanov, Arina N. Pervushina, and Alexey S. Vasilchenko. 2023. ‘Triazoles and Strobilurin Mixture Affects Soil Microbial Community and Incidences of Wheat Diseases’. Plants 12 (3): 660. 10.3390/plants12030660.

[70] Wagg, Cameron, S. Franz Bender, Franco Widmer, and Marcel G. A. van der Heijden. 2014. ‘Soil Biodiversity and Soil Community Composition Determine Ecosystem Multifunctionality’. Proceedings of the National Academy of Sciences 111 (14): 5266–70. 10.1073/pnas.1320054111.

[71] Walder, Florian, Marc W. Schmid, Judith Riedo, Alain Y. Valzano-Held, Samiran Banerjee, Lucie Büchi, Thomas D. Bucheli, and Marcel G. A. van der Heijden. 2022. ‘Soil Microbiome Signatures Are Associated with Pesticide Residues in Arable Landscapes’. Soil Biology and Biochemistry 174 (November):108830. 10.1016/j.soilbio.2022.108830.

[72] Wang, Hongzhe, Wenjie Ren, Yongfeng Xu, Xia Wang, Jun Ma, Yi Sun, Wenbo Hu, et al. 2024. ‘Long-Term Herbicide Residues Affect Soil Multifunctionality and the Soil Microbial Community’. Ecotoxicology and Environmental Safety 283 (September):116783. 10.1016/j.ecoenv.2024.116783.

[73] Wickham, Hadley, Winston Chang, and Maintainer Hadley Wickham. 2016. ‘Package “Ggplot2”‘. Create Elegant Data Visualisations Using the Grammar of Graphics. Version 2 (1): 1–189.

[74] Wickham, Hadley, Romain François, Lionel Henry, Kirill Müller, Davis Vaughan, Posit Software, and PBC. 2023. ‘Dplyr: A Grammar of Data Manipulation’. https://cran.r-project.org/web/packages/dplyr/index.html.

[75] Yachi, Shigeo, and Michel Loreau. 1999. ‘Biodiversity and Ecosystem Productivity in a Fluctuating Environment: The Insurance Hypothesis’. Proceedings of the National Academy of Sciences 96 (4): 1463–68. 10.1073/pnas.96.4.1463.

[76] Zhang, Manyun, Zhihong Xu, Ying Teng, Peter Christie, Jun Wang, Wenjie Ren, Yongming Luo, and Zhengao Li. 2016. ‘Non-Target Effects of Repeated Chlorothalonil Application on Soil Nitrogen Cycling: The Key Functional Gene Study’. Science of The Total Environment 543 (February):636– 43. 10.1016/j.scitotenv.2015.11.053.

[77] Zhang, Ying, Zhao Jiang, Bo Cao, Miao Hu, Zhigang Wang, and Xiaonan Dong. 2011. ‘Metabolic Ability and Gene Characteristics of Arthrobacter Sp. Strain DNS10, the Sole Atrazine-Degrading Strain in a Consortium Isolated from Black Soil’. International Biodeterioration & Biodegradation 65 (8): 1140–44. 10.1016/j.ibiod.2011.08.010.

