## Supporting Information for "Abrupt versus gradual application of pesticides: effects on soil bacterial and fungal communities"

### Supporting Tables

**Table S1:** Pesticide type, family, detected/applied concentration, and mode of action for the ten compounds used in the study.

| Name | Type | Family | Max. monitored concentration (ng a.i. /g soil) | Applied concentration [ng a.i. /g soil] | Mode of Action |
|---|---|---|---|---|---|
| Diflufenican | Herbicide | Carboxamide | 1300 | 1133.3 | It acts as a carotenoid biosynthesis inhibitor, leading to the destruction of chlorophyll, cell rupture, and plant death |
| Napropamide | Herbicide | Alkanamide | 100 | 50.0 | It inhibits the synthesis of certain enzymes in the body so that roots and shoots cannot grow and die (uncertainty of its specific function). |
| Pendimethalin | Herbicide | Dinitroaniline | 1000 | 766.7 | It acts as inhibitor of meristem cell division, suppressing the sprouts and secondary roots of weeds |
| S-metolachlor | Herbicide | Chloroacetamide | 80 | 66.7 | It acts as a cell division inhibitor and mainly inhibits cell growth by inhibiting the synthesis of long-chain fatty acids |
| Boscalid | Fungicide | Carboxamide | 1200 | 1008.3 | It acts as a succinate coenzyme Q reductase inhibitor in the mitochondrial respiratory chain and has a strong inhibitory ability on spore germination |
| Cyproconazole | Fungicide | Triazole | 250 | 204.2 | It acts as a sterol demethylation inhibitor. |
| Epoxiconazole | Fungicide | Triazole | 300 | 237.5 | It acts as a C-14 demethylase inhibitor in sterol biosynthesis, blocking the formation of the cell wall of the bacteria |
| Metrafenone | Fungicide | Benzophenone | 200 | 154.2 | It interferes with the development and formation of appressorium during the germination of pathogenic bacteria and the establishment or formation of polar actin tissue, thereby hindering the normal development and growth of the mycelium. |
| Clothianidin | Insecticide | Neonicotinoid | 60 | 45.8 | It works by combining with nicotinic acetylcholine receptors (nAChRs) of the insect's central nervous system. Immune disrupters can influence in multiple ways the intricate network of interactions among stress agents that have a synergistic impact on the health of insects |
| Imidacloprid | Insecticide | Neonicotinoid | 150 | 133.3 | It acts as a nicotinic acetylcholine receptor agonist, blocking the central nervous system. It disturbs the pest's motor nervous system and causes the failure of chemical signal transmission, paralysis, and death of pests |

**Table S2:** Properties of the environmental fate of the ten pesticides used in the study.

| Name | Type | Family | Log($K_{OW}$) | Log($K_{AW}$) | $DT_{50}$ | $K_{foc}$ | GUS | BCF |
|---|---|---|---|---|---|---|---|---|
| Diflufenican | Herbicide | Carboxamide | 4.2 | -1.9 | 94.5 | 2215 | 1.19 | 1276 |
| Napropamide | Herbicide | Alkanamide | 3.3 | -4.1 | 70 | 885 | 1.96 | 98 |
| Pendimethalin | Herbicide | Dinitroaniline | 5.4 | 0.10 | 182.3 | 13792 | -0.28 | 5100 |
| S-metolachlor | Herbicide | Chloroacetamide | 3.0 | -2.7 | 51.8 | 200.2 | 2.32 | 68.8 |
| Boscalid | Fungicide | Carboxamide | 3.0 | -4.3 | 484.4 | 772 | 2.68 | 107 |
| Cyproconazole | Fungicide | Triazole | 3.1 | -4.3 | 142 | 364 | 3.04 | 28 |
| Epoxiconazole | Fungicide | Triazole | 3.3 | -4.8 | 353.5 | 894 | 2.09 | 70 |
| Metrafenone | Fungicide | Benzophenone | 4.3 | -0.88 | 200.9 | 3105 | 0.91 | 530 |
| Clothianidin | Insecticide | Neonicotinoid | 0.91 | -10.5 | 545 | 160 | 3.74 | 0 |
| Imidacloprid | Insecticide | Neonicotinoid | 0.57 | -9.8 | 191 | 225 | 3.69 | 0.61 |

Log($K_{OW}$): < 2.7 = Low bioaccumulation, 2.7 – 3 = Moderate, > 3.0 = High
Log($K_{AW}$): > 100 = Volatile, 0.1 - 100 = Moderately volatile, < 0.1 = Non-volatile
$DT_{50}$: < 30 = Non-persistent, 30 - 100 = Moderately persistent, 100 - 365 = Persistent, > 365 = Very persistent
$K_{oc}$/ $K_{foc}$: < 15 = Very mobile, 15 - 75 = Mobile, 75 - 500 = Moderately mobile
GUS: > 2.8 = High leachability, 2.8 - 1.8 = Transition state, < 1.8 = Low leachability
BCF: < 100 = Low potential, 5000 – 100 = Threshold for concern, > 5000 – High potential

### Supporting Results

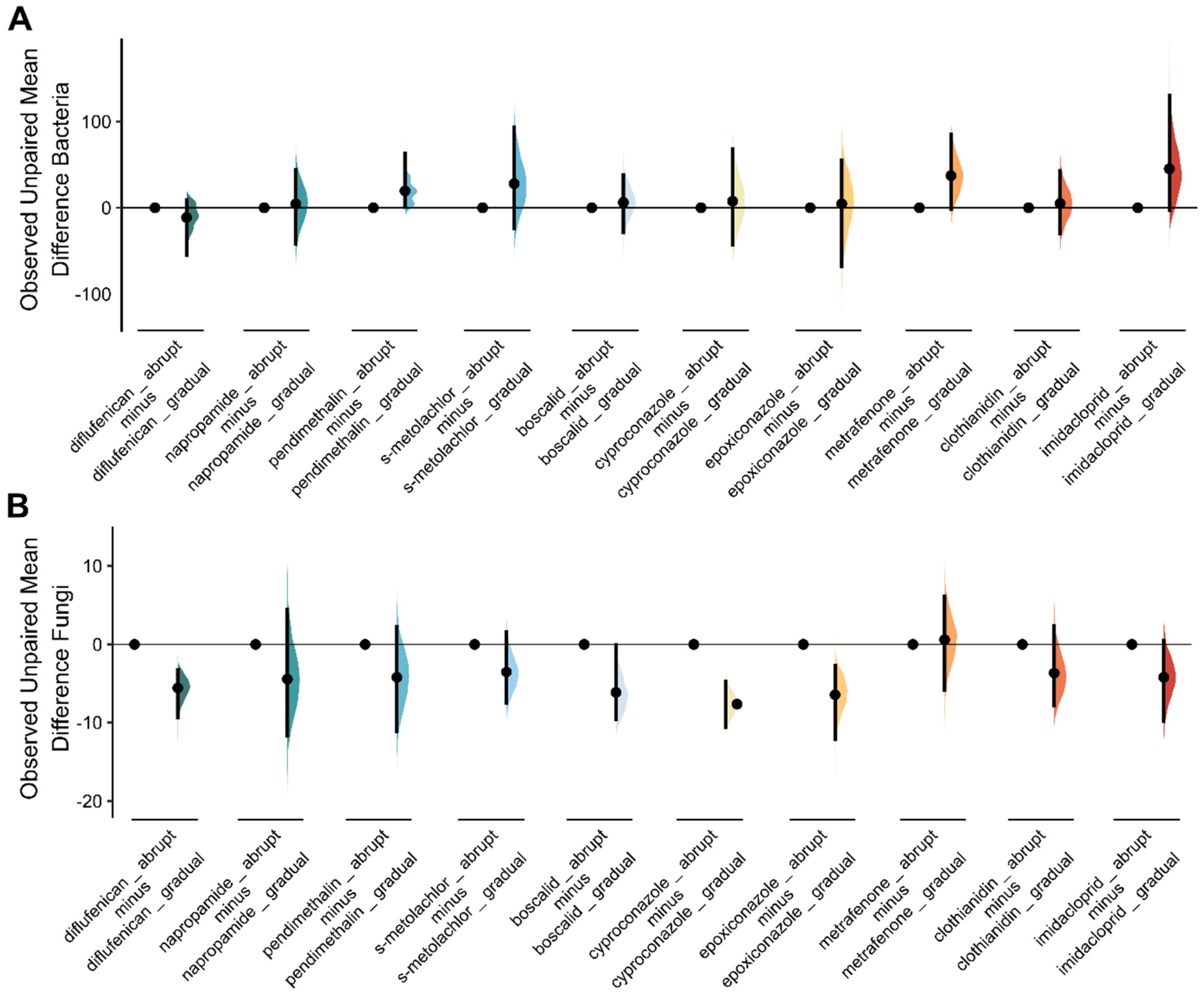

**Figure S1:** Mean difference in bacterial (A) and fungal (B) inverse Simpson's dominance effect sizes of individual treatments, with circles representing the bootstrapped effect size, mean (effect magnitude), and vertical lines corresponding to the 95% confidence interval (effect precision). The density plots show the bootstrapped data distribution.

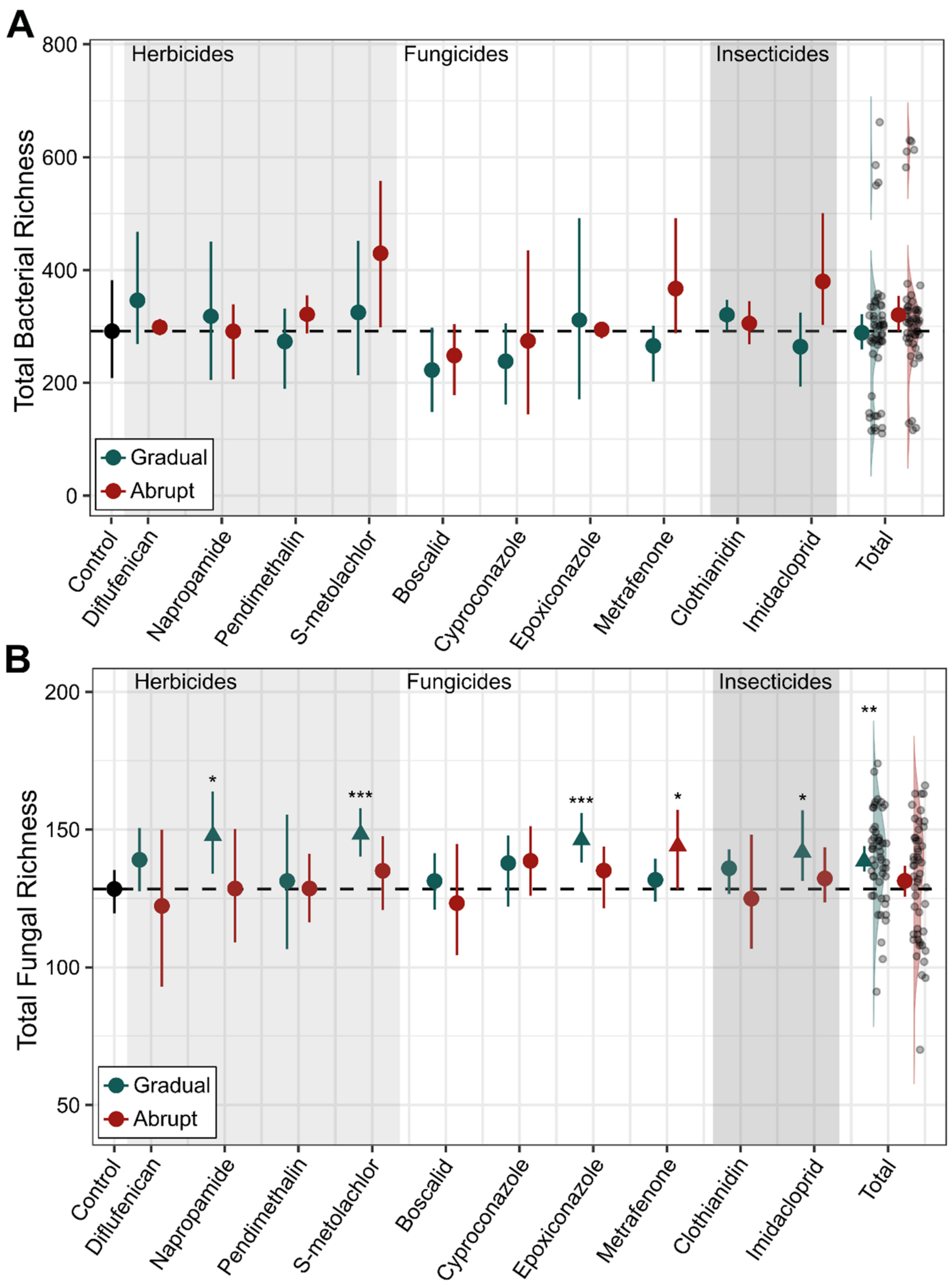

**Figure S2:** Individual effects of different pesticides (ordered by herbicides, fungicides, and insecticides) applied gradually (blue) or abruptly (red) on total richness. The circles and triangles, along with the error bars, represent the mean effect size and 95% confidence intervals (CIs), respectively. The circles represent neutral effects, while the triangles (with an arrow pointing up or down) represent significant positive or negative effects. The asterisks mark the degree of significance (*: P-value < 0.05, **: P-value < 0.01, ***: P-Value < 0.001). For bacteria (A), no significant change in bacterial richness was observed for any of the ten pesticides. For fungi (B), a significant increase in fungal richness was observed for four of the ten pesticides in the gradual treatment group, while only metrafenone showed a significant increase in the abrupt treatment group. In addition, the overall gradual treatment was found to be significant.

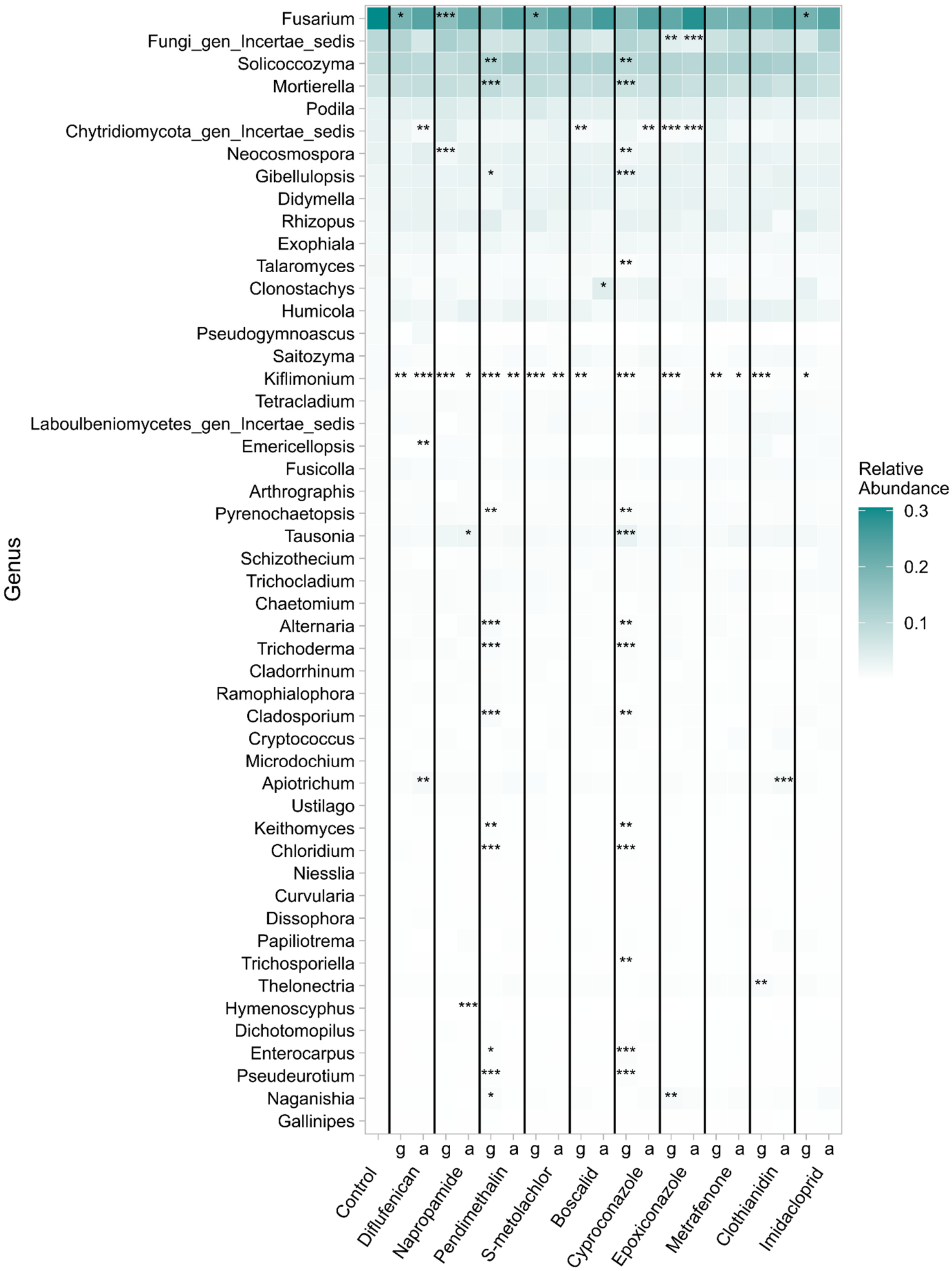

**Figure S3:** Heatmap showing the change in mean relative abundance of fungal genera as a function of pesticide application (g = gradual; a = abrupt) for the 50 most abundant genera, sorted by highest to lowest abundance. Red marks the decrease in abundance compared to the control, and blue marks the increase. The asterisks denote the degree of significance (P-value < 0.05, **: P-value < 0.01, ***: P-value < 0.001) in change of abundance compared to the control.

**Table S3:** Summary of PERMANOVA and dispersion test results comparing microbial community composition among treatment groups. The overall PERMANOVA results (top section for each pesticide) include degrees of freedom (Df), sum of squares (SumSq), F-statistic (F_Model), $R^2$ values, and overall model P-value. The pairwise PERMANOVA results (bottom section for each pesticide) present comparisons between treatment groups, including $R^2$, F-statistic, P-values, and dispersion test P-values.

| Diflufenican | Df | SumSq | F_Model | R2 | P_value | |
|---|---|---|---|---|---|---|
| Model | 2 | 0.359 | 3.330 | 0.281 | 0.001 | |
| Residual | 17 | 0.917 | NA | 0.719 | NA | |
| Total | 19 | 1.277 | NA | 1.000 | NA | |
| Dispersion | 2 | 0.005 | 0.528 | NA | 0.599 | |
| | Df | SumSq | F_Model | R2 | P_value | Dispersion |
| Control vs gradual | 1 | 0.248 | 4.750 | 0.268 | 0.002 | 0.354 |
| Control vs abrupt | 1 | 0.187 | 3.231 | 0.199 | 0.002 | 0.915 |
| Gradual vs abrupt | 1 | 0.066 | 1.314 | 0.141 | 0.122 | 0.336 |

| Napropamide | Df | SumSq | F_Model | R2 | P_value | |
|---|---|---|---|---|---|---|
| Model | 2 | 0.508 | 3.894 | 0.314 | 0.001 | |
| Residual | 17 | 1.110 | NA | 0.686 | NA | |
| Total | 19 | 1.618 | NA | 1.000 | NA | |
| Dispersion | 2 | 0.003 | 0.311 | NA | 0.737 | |
| | Df | SumSq | F_Model | R2 | P_value | Dispersion |
| Control vs gradual | 1 | 0.370 | 5.758 | 0.307 | 0.001 | 0.446 |
| Control vs abrupt | 1 | 0.241 | 3.985 | 0.235 | 0.004 | 0.738 |
| Gradual vs abrupt | 1 | 0.100 | 1.336 | 0.143 | 0.149 | 0.676 |

| Pendimethalin | Df | SumSq | F_Model | R2 | P_value | |
|---|---|---|---|---|---|---|
| Model | 2 | 0.466 | 3.349 | 0.283 | 0.001 | |
| Residual | 17 | 1.184 | NA | 0.717 | NA | |
| Total | 19 | 1.650 | NA | 1.000 | NA | |
| Dispersion | 2 | 0.024 | 1.186 | NA | 0.329 | |
| | Df | SumSq | F_Model | R2 | P_value | Dispersion |
| Control vs gradual | 1 | 0.329 | 4.266 | 0.247 | 0.001 | 0.304 |
| Control vs abrupt | 1 | 0.190 | 3.563 | 0.215 | 0.003 | 0.493 |
| Gradual vs abrupt | 1 | 0.154 | 1.835 | 0.187 | 0.016 | 0.263 |

| Cyproconazole | Df | SumSq | F_Model | R2 | P_value | |
|---|---|---|---|---|---|---|
| Model | 2 | 0.431 | 2.897 | 0.254 | 0.001 | |
| Residual | 17 | 1.265 | NA | 0.746 | NA | |
| Total | 19 | 1.696 | NA | 1.000 | NA | |
| Dispersion | 2 | 0.028 | 0.964 | NA | 0.401 | |
| | Df | SumSq | F_Model | R2 | P_value | Dispersion |
| Control vs gradual | 1 | 0.342 | 4.119 | 0.241 | 0.002 | 0.344 |
| Control vs abrupt | 1 | 0.146 | 2.708 | 0.172 | 0.002 | 0.497 |
| Gradual vs abrupt | 1 | 0.131 | 1.395 | 0.148 | 0.026 | 0.392 |

| Epoxiconazole | Df | SumSq | F_Model | R2 | P_value | |
|---|---|---|---|---|---|---|
| Model | 2 | 0.264 | 2.514 | 0.228 | 0.001 | |
| Residual | 17 | 0.894 | NA | 0.772 | NA | |
| Total | 19 | 1.158 | NA | 1.000 | NA | |
| Dispersion | 2 | 0.004 | 0.476 | NA | 0.629 | |
| | Df | SumSq | F_Model | R2 | P_value | Dispersion |
| Control vs gradual | 1 | 0.229 | 4.129 | 0.241 | 0.001 | 0.691 |
| Control vs abrupt | 1 | 0.069 | 1.298 | 0.091 | 0.161 | 0.444 |
| Gradual vs abrupt | 1 | 0.083 | 1.740 | 0.179 | 0.077 | 0.480 |

| Metrafenone | Df | SumSq | F_Model | R2 | P_value | |
|---|---|---|---|---|---|---|
| Model | 2 | 0.412 | 3.595 | 0.297 | 0.001 | |
| Residual | 17 | 0.975 | NA | 0.703 | NA | |
| Total | 19 | 1.387 | NA | 1.000 | NA | |
| Dispersion | 2 | 0.002 | 0.251 | NA | 0.781 | |
| | Df | SumSq | F_Model | R2 | P_value | Dispersion |
| Control vs gradual | 1 | 0.270 | 4.552 | 0.259 | 0.001 | 0.822 |
| Control vs abrupt | 1 | 0.242 | 4.398 | 0.253 | 0.001 | 0.650 |
| Gradual vs abrupt | 1 | 0.056 | 0.966 | 0.108 | 0.471 | 0.335 |

**Table S3:** continued

| S-metolachlor | Df | SumSq | F_Model | R2 | P_value | |
|---|---|---|---|---|---|---|
| Model | 2 | 0.343 | 3.259 | 0.277 | 0.001 | |
| Residual | 17 | 0.895 | NA | 0.723 | NA | |
| Total | 19 | 1.238 | NA | 1.000 | NA | |
| Dispersion | 2 | 0.010 | 1.201 | NA | 0.325 | |
| | **Df** | **SumSq** | **F_Model** | **R2** | **P_value** | **Dispersion** |
| Control vs gradual | 1 | 0.268 | 4.602 | 0.261 | 0.002 | 0.981 |
| Control vs abrupt | 1 | 0.133 | 2.664 | 0.170 | 0.001 | 0.151 |
| Gradual vs abrupt | 1 | 0.084 | 1.762 | 0.181 | 0.055 | 0.142 |

| Clothianidin | Df | SumSq | F_Model | R2 | P_value | |
|---|---|---|---|---|---|---|
| Model | 2 | 0.414 | 3.518 | 0.293 | 0.001 | |
| Residual | 17 | 1.001 | NA | 0.707 | NA | |
| Total | 19 | 1.415 | NA | 1.000 | NA | |
| Dispersion | 2 | 0.000 | 0.055 | NA | 0.947 | |
| | **Df** | **SumSq** | **F_Model** | **R2** | **P_value** | **Dispersion** |
| Control vs gradual | 1 | 0.292 | 5.128 | 0.283 | 0.003 | 0.898 |
| Control vs abrupt | 1 | 0.195 | 3.272 | 0.201 | 0.003 | 0.839 |
| Gradual vs abrupt | 1 | 0.098 | 1.612 | 0.168 | 0.090 | 0.619 |

| Boscalid | Df | SumSq | F_Model | R2 | P_value | |
|---|---|---|---|---|---|---|
| Model | 2 | 0.303 | 2.617 | 0.235 | 0.002 | |
| Residual | 17 | 0.983 | NA | 0.765 | NA | |
| Total | 19 | 1.286 | NA | 1.000 | NA | |
| Dispersion | 2 | 0.000 | 0.020 | NA | 0.980 | |
| | **Df** | **SumSq** | **F_Model** | **R2** | **P_value** | **Dispersion** |
| Control vs gradual | 1 | 0.234 | 4.038 | 0.237 | 0.001 | 0.963 |
| Control vs abrupt | 1 | 0.112 | 1.951 | 0.130 | 0.033 | 0.837 |
| Gradual vs abrupt | 1 | 0.088 | 1.488 | 0.157 | 0.126 | 0.855 |

| Imidacloprid | Df | SumSq | F_Model | R2 | P_value | |
|---|---|---|---|---|---|---|
| Model | 2 | 0.359 | 3.176 | 0.272 | 0.001 | |
| Residual | 17 | 0.960 | NA | 0.728 | NA | |
| Total | 19 | 1.319 | NA | 1.000 | NA | |
| Dispersion | 2 | 0.001 | 0.063 | NA | 0.939 | |
| | **Df** | **SumSq** | **F_Model** | **R2** | **P_value** | **Dispersion** |
| Control vs gradual | 1 | 0.290 | 5.197 | 0.286 | 0.001 | 0.758 |
| Control vs abrupt | 1 | 0.123 | 2.134 | 0.141 | 0.021 | 0.893 |
| Gradual vs abrupt | 1 | 0.099 | 1.767 | 0.181 | 0.023 | 0.790 |
